# EPITHELIAL REMODELING AND MICROBIAL DYSBIOSIS IN THE LOWER RESPIRATORY TRACT OF VITAMIN A-DEFICIENT MOUSE LUNGS

**DOI:** 10.1101/2024.06.21.600110

**Authors:** Kiloni R. Quiles, Feng-Zhi Shao, W. Evan Johnson, Felicia Chen

## Abstract

The World Health Organization identified vitamin A deficiency (VAD) as a major public health issue in low-income communities and developing countries, while additional studies have shown dietary VAD leads to various lung pathologies. Once believed to be sterile, research now shows that transient microbial communities exist within healthy lungs and are often dysregulated in patients suffering from malnourishment, respiratory infections, and disease. The inability to parse vitamin A-mediated mechanisms from other metabolic mechanisms in humans with pathogenic endotypes, as well as the lack of data investigating how VAD affects the lung microbiome, remains a significant gap in the field. To address this unmet need, we compared molecular, metatranscriptomic, and morphometric data to identify how dietary VAD affects the lung as well as the lung microbiome. Our research shows structural and functional alterations in host-microbe-diet interactions in VAD lungs compared to vitamin A-sufficient (VAS) lungs; these changes are associated with epithelial remodeling, a breakdown in mucociliary clearance, microbial imbalance, and altered microbial colonization patterns after 8 weeks of vitamin A deficient diet. These findings confirm vitamin A is critical for lung homeostasis and provide mechanistic insights that could be valuable for the prevention of respiratory infections and disease.

**ONE SENTENCE SUMMARY:** Vitamin A deficiency leads to epithelial remodeling, ciliary dysfunction, a loss of mucociliary clearance, and an increase in biofilm formation and microbial abundance in adult mouse lungs.

## INTRODUCTION

### Vitamin A deficiency and host lung pathology

Vitamin A deficiency (VAD), combined with protein malnutrition, is the most common nutritional disorder in the world.^1^ According to the World Health Organization (WHO), VAD primarily affects low-income communities and developing countries, two demographics where recurrent respiratory infections are common and malnutrition is widespread.^1,2^ It is well-documented that vitamin A plays an essential role in the maintenance of mucociliary differentiation of the tracheobronchial tree in vertebrates, including humans.^3–5^ Animals deficient in vitamin A display respiratory epithelial hyperplasia followed by squamous metaplasia.^6^ Similarly, airway epithelial culture in-vitro in serum free of retinoic acid (RA), the bioactive metabolite of vitamin A, loses the pseudostratified mucociliary epithelium and possesses a stratified, squamous morphology.^7^ The loss of pseudostratified mucociliary epithelium is a hallmark of several respiratory diseases characterized by epithelial injuries, repair defects, tissue remodeling, and altered mucociliary clearance (MCC).^8,9^ Indeed, chronic VAD in humans predisposes the host to COPD, asthma, pulmonary fibrosis, and respiratory infections such as TB, influenza, measles, and SARS-CoV-2).^10–16^

While these findings are noteworthy, determining causality with respect to diet-associated disease progression in human samples is difficult. Patients deficient in one vitamin are often deficient in other micronutrients.^17^ Combined with other cofactors such as lifestyle and genetic background, human studies lack the specificity needed to characterize the role of dietary vitamin A in lung function and homeostasis. A well-controlled mouse model can overcome these challenges. By administering a custom diet to isogenic littermates, we can isolate the functional and morphological effects of dietary VAD on the lung without undue nutritional or genetic influence.

Another major clinical challenge is the inability to fully reverse the epithelial damage, risk of infection, and rate of mortality with vitamin A supplementation alone.^18,19^ In some cases, malnourished patients given excess vitamin A can cause a cytotoxic condition known as hypervitaminosis A in patients.^12^ leading to symptom exacerbation and a worsened prognosis overall.^20^ Because of this limitation, investigating secondary mechanisms implicated in the pulmonary-vitamin A pathway is critical to fully characterize how vitamin A regulates lung homeostasis. Conversely, respiratory infections have been shown to deplete vitamin A serum levels and retinoid stores in the liver of patients, effectively inducing VAD independent of diet.^21^ Thus, identifying alternative therapeutic targets would not only impact the long-term health of patients, but may also be an effective prevention and control strategy for the transmission of respiratory infections to the greater public.

### The lung and the microbiome

It has been well-characterized that infectious agents impair lung function and repair mechanisms in VAD patients, but these studies fail to account for more complex host-microbe and microbe-microbe interactions, leaving the mechanisms involved in these interactions largely unknown.^22^ Additionally, there are associations between the composition and diversity of the lower respiratory tract (LRT) microbiome and idiopathic pulmonary fibrosis (IPF), asthma,^23^ ARDS,^24^ COPD,^25^ lung cancer,^26^ and SARS-CoV-2-induced lung damage.^27^ These findings suggest that the LRT microbial microenvironment may trigger, predispose, or exacerbate lung pathogenesis.

The airways are lined by a pseudostratified layer of epithelial cells which plays a major role in microbial elimination. Secretory cells secrete a mucinous layer that protects the apices of the airway epithelium by trapping the inhaled particles, potential pathogens, and commensal microbes within the mucin. Ciliary cells work in concert to expel these substances through a process called the *mucociliary escalator.*^9^ The cilia beat in a semi-synchronous fashion to move the mucin-trapped microbes up and out of the lung; in this way, the lung is a unique microenvironment due to the transient nature of its microbial communities. When mucociliary clearance (MCC) is impaired, the microbes cannot be expelled effectively, providing a permissive niche for pathogenic microbial colonization and biofilm formation;^28^ the subsequent dysbiosis may cause further epithelial injury, leading to chronic diseases. Here, we use a combined metatranscriptomic, morphometric, and functional approach to explore changes to the host lung epithelium as well as characterize the microbial milieu of vitamin A-deficient (VAD) mouse lungs.

## RESULTS

### ESTABLISHMENT OF THE VITAMIN A-DEFICIENT MOUSE MODEL

Studies have shown serum and liver vitamin A levels do not accurately reflect the bioavailability of RA in the tissue^12^. This is because the concentration of RA is tightly regulated at both the tissue and cellular levels. Dietary vitamin A is absorbed into the small intestine and circulates in the body as retinol, where it is then taken into the cells. When the tissue has sufficient or excess RA, the retinol is converted to retinyl ester for storage, a process mediated by lecithin retinol acyltransferase (LRAT).^29^ However, when there is a deficiency of RA (RAD), the oxidation of retinol to RA occurs, with a key step mediated by aldehyde dehydrogenase (ALDH).^30^ RA then enters the nucleus where it binds to retinoid X receptor (RXR) and retinoic acid receptor (RAR) heterodimer known as the retinoic acid response element (RARE) to mediate transcriptional activity (Figure 1B).^31–32^

**Fig. 1.**
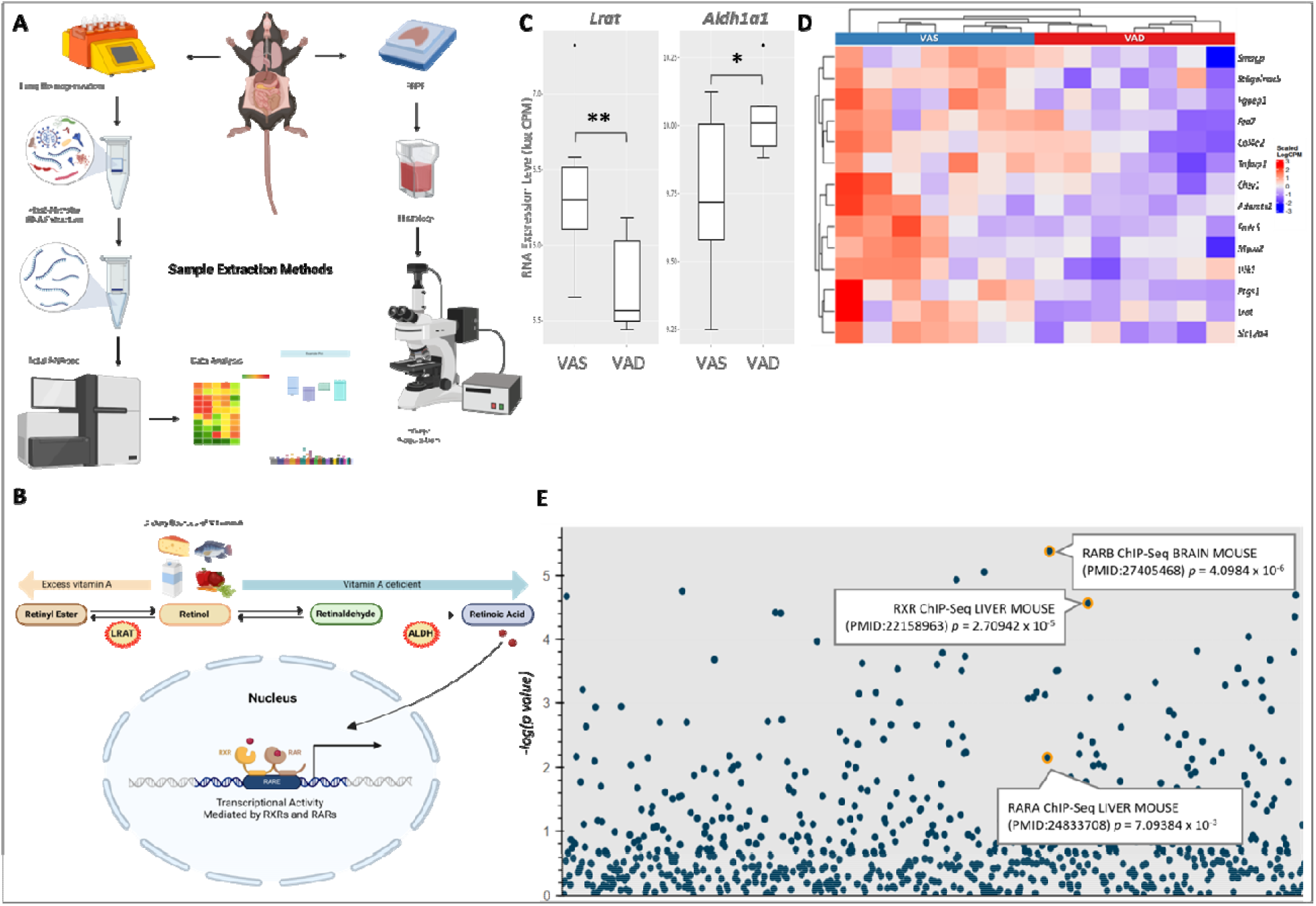
Transcriptional confirmation of vitamin A deficiency In adult mouee lunge after 8 weeks of detery modulation. (A) General schematic of the mixed host-mlcroblome RNA extraction and data analysis protocol. (B) Metabolic pathway of Vitamin A. (C, Gene expression of *Lret* and *A)dh1a1,* two Indicators of tissue vitamin A status. *P-veluaa* = 0.009 and 0.038 respectively. (D) Heatmap of vitamin A targets from literature In Vitamin Arsufflclent (VAS) and vitem n A-deflclent (VAD, mouse lungs. (E) Manhattan plot of ChlR-X Enrichment Analysis (СҺЕА) gene sals. Each point Ind cates one term from the СҺЕА 2022 gene set. The y-axls represents the -log(p-velue) of the enrichment between tne Input gene set and the СҺЕА term gene set. The Input gene set used Included ell downreguleted genes In VAS lungs at 8 weeks with a p value < 0.0S (n—14). All mouse RNA sequencing tiles were annotated and processed using SlngleCell Toolkit version 1.9.1 on Rstudlo version 4.1.1. Figure B created using Blorender.

To determine whether we can achieve vitamin A deficiency in the adult mouse lung with dietary modulation, we maintained 8-week old wild-type C57BL/6J mice on a VAS or VAD diet for 8 weeks. We extracted the left lobes of the lung for bulk RNA sequencing while the right lobes were embedded for histology and immunohistochemistry. The sequences were then trimmed, aligned, and processed for downstream analysis (Figure 1A).

Decreased *Lrat* expression indicates the conversion of retinol to the reversible storage form retinyl ester is reduced in VAD lungs. Conversely, increased *Aldh1a1* expression indicates the rate of conversion from retinol to retinoic acid (RA) has also increased (Figure 1C).

Differential expression of genes reported in literature to be associated with human VAD status accurately clustered our VAS and VAD samples (Figure 1D).^33–40^ Finally, genes downregulated in VAD compared to VAS lungs (*p <* 0.05) were analyzed against gene sets from the ChEA transcription factor target database.^41^ Gene sets associated with retinoic/retinoid receptors were found highly enriched in our samples (Figure 1E).^42^ These findings confirm that tissue-specific VAD is achieved in wild-type mice after removing vitamin A from their diet for 8 weeks.

### VAD LEADS TO EPITHELIAL REMODELING IN ADULT MOUSE LUNGS

After 8 weeks of dietary vitamin A deficiency, we stained whole lung samples to analyze the morphological changes to the proximal and distal airways of the lower respiratory tract (LRT), the region of the lung below the branching of the trachea into the primary bronchi (Figures 2A-B). Our results show epithelial remodeling in the airways (Figures 2C-F) of VAD mouse lungs (right) compared to the VAS controls (left). The observed VAD-induced airway remodeling is characterized by epithelial denudation and cellular debris, focal hyperplasia, thickening of the basement membrane, and an increase in protein expression of proliferative marker Ki-67 in the airway epithelium of both the proximal (Figure 2G-H) and distal regions (Figures 2I-J) of the LRT.^43^ These findings indicate an increase in proliferation of airway epithelial cells in VAD lungs.

**Fig. 2.**
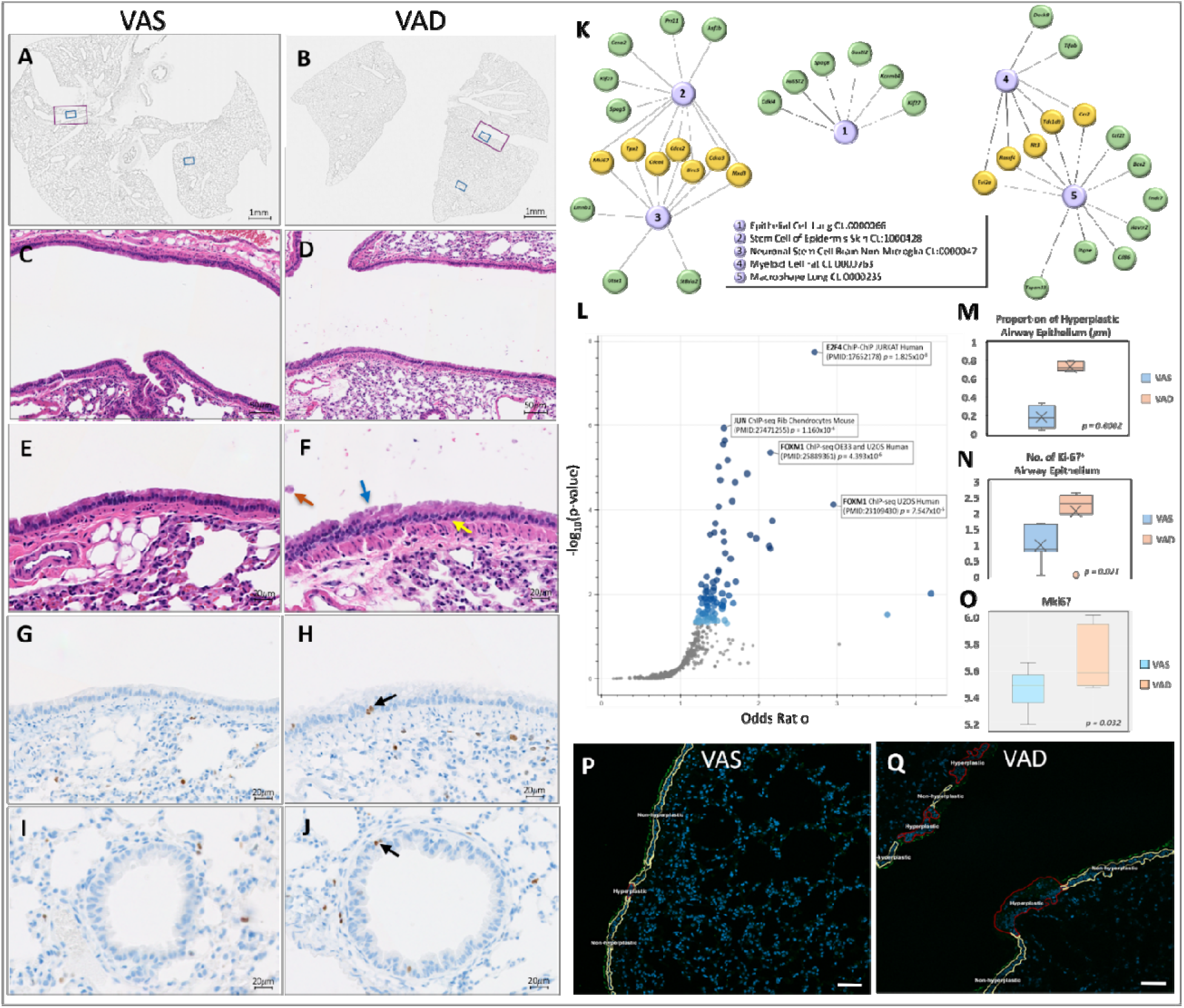
Increased proliferation and epithelial remodeling observed in VAD airways after 8 weeks. (A-J) Histology staining of VAS and VAD lungs. (A,B) Whole lung scans showing regions of image analysis in the proximal (E-H) and distal (I,J) regions of the LRT. Hematoxylin and Eosin (C-F) and Ki-67 (G-J) staining of VAS (left) and VAD (right) lungs. Hyperplasia in the airway is indicated by the blue arrow. Epithelial denudation is indicated by the orange arrow. Thickening of the basement membrane is indicated by the yellow arrow. Ki-67 positive airway epithelium are indicated by the black arrows. Photos taken at 50um (C-D) and 20um (E-J) magnification. (K) Enrichr KG cluster map depicting five major nodes and three major clusters of gene signatures from the Tabula Muris mouse model transcriptome database. The input gene set used included all upregulated genes in VAD lungs at 8 weeks with a p value < 0.05 (n= 14). (L) Volcano plot showing targets of pro-proliferation transcriptional factors FOXM1, JUN, and E2F4 enriched in VAD lungs at 8 weeks (p < 0.05, n=14). (M) Proportion of hyperplastic epithelium in VAD airways compared to controls at Ө weeks. Boxplots of (N) KI67-posltlve cell counts In the airways and (O) Mkl67 RNA expression In VAS and VAD lungs at 8 weeks (n=14). Hyperplastic ratios were quantified using QuPath (P,Q). Scale bar = 50um.

To determine if the transcriptional signature implicates airway epithelium as the major cell types affected by host VAD status, we compared the genes upregulated in VAD lungs (p < 0.05) with gene signatures from the Tabula Muris mouse model transcriptome single-cell database.^44^ Lung epithelium, ectodermal epithelial stem cells, and myeloid cells (Figure 2K) as well as pro-proliferation transcriptional factors FOXM1, JUN, and E2F4 (Figure 2L)^45–47^ were enriched in our VAD dataset, suggesting a measurable increase in epithelial and immune cell signatures following dietary modulation.

Quantitative analysis shows a marked increase in focal regions of hyperplastic epithelium in VAD airways (Figure 2M, p = 0.0002). Further, quantification of Ki67-positive stained airway epithelial cells (Figure 2N) and the expression level of *Mki67*, the transcript for Ki-67 protein, (Figure 2O) both support increased proliferation in VAD airway epithelium (Figures 2P-Q, outlined in red). These results were consistent with previously published data showing that vitamin A/RA pathway inhibition promotes proliferation in mouse and human lung epithelial organoids.^48^

### MUCOCILIARY CLEARANCE IS IMPAIRED IN VAD LUNGS

To investigate which cell types are implicated in the observed VAD-induced remodeling of the airway epithelium, we first looked at the RNA expression of the major epithelial cell types found in the LRT. We measured secretory cell markers SCGB1A1^48^ and SCGB3A2^49^ and ciliated cell markers FOXJ1^50^ and TUBB4A^51^ (Figure 3A) and found a significant increase in the gene expression of secretory cells in VAD lungs. Using images stained with histology and immunohistochemistry, we quantified secretory and multiciliated epithelial cell proportions in the airways (Figure 3B). These results also showed an increase in secretory epithelium in VAD airways compared to VAS, indicating that dietary VAD-induced focal hyperplasia increases the proportion of secretory cells within the airways of the LRT.

**Fig. 3.**
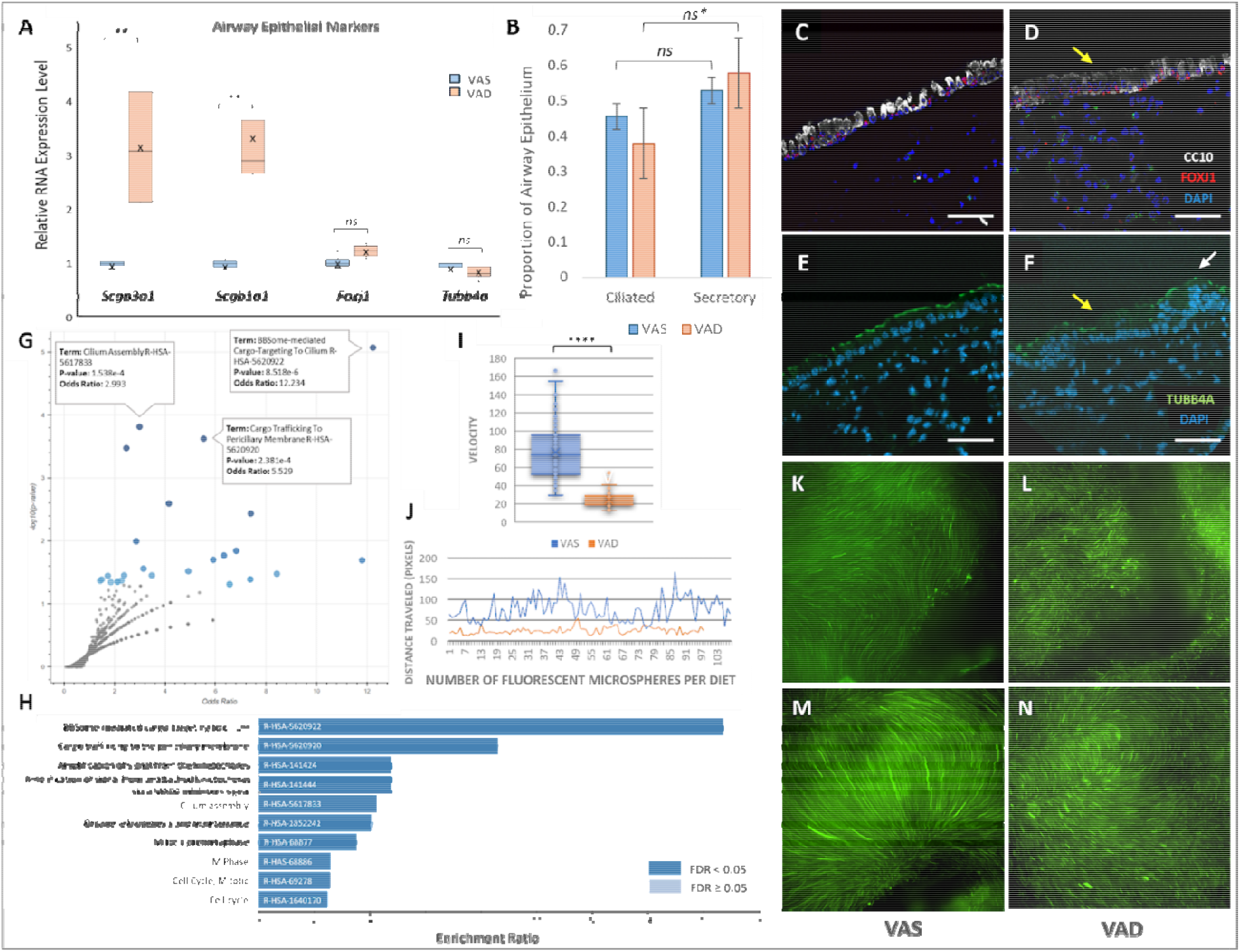
VAD-induced airway remodeling leads to a breakdown in mucociliary clearance. (A) Gene expression of secretory *(Scgblal* and *ScgbSaS)* and diary (Feutff, *Tubb4a}* markers In VAS and VAD mouse lungs (n—14). ** ▪ p < 0.01. *“na”* ▪ not significant. (B) Proportion of ciliated and secretory cells within the airway of the LRT In 8-weak VAS and VAD lungs. The y-axls represents the actual proportion of total airway epithelium where 1 = 100%. P-values = 0.26 (VAS) and 0.07 (VAD) respectively, ne* = near significance. (C-F) Immunohistochemical staining of airway epithelium. Secretory cells were visualized using CC10 while dilated cells were visualized using both nuclear FoxJ1 and cllla-speclflc Tubb4a. Yellow arrows ndlcete regions of reduced apical proton express on. White arrows Indicate focal hypeiplaala. (G) Volcano plot of the top pathways altered In VAD lungs, analyzed using the Reactome 2022 database. The Input dataset Included all dlfferentlally expressed genes (DEGs) with p < 0.06 (n^ 19). (H) Barplot showing the top 10 enriched pathways from the Enrlchr Reactome database using only genes upregulated In VAD lungs with p < 0.05. (kJ) Fluorescent microsphere velocity (I) and overall distance traveled (J) In VAS and VAD lungs, p < 0.00001, t-valua = 17.70024. (K-N) Composte mages of fluorescent mlcrospnere motility In the trachea of VAS and VAD lungs from a cross-sectional (K-L) and anterior (M-N) view at 20x magnification. All samples represented In mis figure were collected at me 8-waek tlmepokit

We next performed immunohistochemistry to visualize the expression of secretory epithelial cell marker CC10 (the protein product of the transcript *Scgb1a1*), ciliated nuclear marker FOXJ1 (Figures 3C-D) and apical ciliary marker Tubb4a (Figures 3E-F).^50–52^ Briefly, several studies have found secretory protein CC10 expression upregulated in obstructive lung diseases and downregulated in restrictive lung diseases, making CC10 expression an important biomarker for lung health and disease.^53^ Likewise, expression of nuclear FOXJ1 and ciliated TUBB4A proteins are known key regulators of motile ciliogenesis in the airway epithelium.^54^ SCGB1A1, a gene typically expressed within the secretory granules of club cells located at the apical portion of the cytoplasm, is now observed throughout the cytoplasm and along the basolateral region of club cells in hyperplastic regions of the VAD airway (Figures 3C-D). Furthermore, immunostaining of TUBB4A expression is normally restricted to the cilia on the apical surface of multiciliated epithelium under VAS conditions. Our data demonstrates significantly reduced expression in the hyperplastic region of the VAD airway (Figures 3E-F). These findings confirm that dietary VAD leads to phenotypic changes characterized by a loss of apical protein expression of both major cells types in the intraparenchymal airways, two proteins that are critical for maintaining lung homeostasis through mucociliary clearance mechanisms.^55^

To determine the gene signature associated with these phenotypic changes, we performed gene set enrichment analysis of all differentially expressed genes in VAD lungs at 8 weeks (n=14, 619 genes with p < 0.05). The top three significant pathways from the Reactome 2022 database that were enriched in our VAD dataset were associated with the cilia assembly and trafficking (Figures 3G-H). Of note, these same pathways were enriched whether we used a gene set with all differentially expressed genes (DEGs) from VAD lungs (Figure 3G) or if we restricted our input gene set to only upregulated genes in the VAD lung (Figure 3H). These findings provide robust evidence of a cilia-specific gene signature present in VAD lungs. Interestingly, the upregulated VAD gene set also showed enrichment for cell cycle pathways such as “Mitotic prometaphase”, “M phase”, “Cell cycle, mitotic”, and “Cell cycle” (Figure 3H), providing further evidence that dietary VAD leads to a hyperproliferative and ciliopathic gene signature in the LRT.

Multiciliated cell structure and function is imperative for proper mucociliary clearance (MCC), the primary innate defense mechanism in the airway.^56^ To determine how these morphologic, genetic, and transcriptional changes affect MCC, we then performed an ex-vivo ciliary motility assay to visualize the cilia-driven flow in VAS and VAD airways.^57^ Briefly, tracheal explants were extracted and immediately placed in petri dishes containing L15 media with 20_µ_l/ml of 0.2um fluorescent microspheres; this microsphere size is representative of most small particles, microbes, and other pollutants our lungs commonly trap and expel under homeostatic conditions. Each petri dish was recorded using a Nikon Deconvolution Wide-Field Fluorescence Microscope (B15D) equipped with a camera and video acquisition software to measure the cilia-driven flow at 200 frames per second. Video frames were concatenated to create still images, allowing us to measure the velocity and distance traveled of individual beads (green lines). A minimum of 100 beads were measured for each arm of the study.

In VAS lungs, the fluorescent beads are observed moving in a smoother, more efficient motion and follow a largely singular direction, indicating that cilia-driven propulsion in the VAS trachea is intact (Figures K,M). Conversely, VAD tracheas showed a significant reduction in velocity (Figure 3I, p<0.00001) and distance traveled (Figure 3J) of fluorescent microspheres over the same period of time (Figures 3L,N). Taken together, these findings confirm that dietary VAD leads to epithelial remodeling characteristic of ciliary dysfunction in the intraparenchymal airways, causing a breakdown in mucociliary clearance.

### GENE EXPRESSION ANALYSIS OF THE LUNG MICROBIOME INDICATES DYSBIOSIS AND PREDISPOSITION TO DISEASE IN VAD LUNGS

Ciliary dysfunction leads to MCC impairment, one of the most important mechanisms for maintaining proper lung homeostasis.^58^ Ciliary dysfunction is commonly found in patients with Primary Ciliary Dyskinesia (PCD) and cystic fibrosis (CF), but it can also be observed in patients with α1-antitrypsin deficiency (AATD), chronic obstructive pulmonary disease (COPD), and asthma.^59^ And while research has uncovered evidence suggesting diet influences the severity and progression of pulmonary pathogenesis, the exact etiologies that influence prognosis and mortality risk in these patients remain largely unknown. To date, there have been few longitudinal studies focusing on compositional changes to the lung microbiome over the course of a disease, and even fewer investigating how diet directly affects lung health over time. Longitudinal microbiome data from humans also tend to contain large batch effects due to the variability in cohorts, geographic location, extraction techniques, patient involvement, and environmental exposures.^60^ Luckily, the function and structure of cilia-related proteins responsible for proper motility and MCC are evolutionarily conserved, and can be easily studied in relation to the human lung using a standard isogenic mouse model.^61^ Based on our findings, we propose that chronic VAD may predispose adult mouse lungs to respiratory infections over time.

For the purposes of this study, we define “chronic VAD” as maintaining an adult mouse on a VAD diet for 8 weeks, which is the equivalent of a human eating a VAD diet for approximately 7 years.^62^ We extracted the microbial RNA from whole lung homogenates to sequence and analyze for compositional changes to the microbiome after 3 weeks and 8 weeks of dietary modulation respectively. Alpha diversity, or the measurement of diversity *within* samples^63^ (Figure 4A), was performed on bacterial read counts at the Phylum level using the Mann-Whitney/Kruskal-Wallis statistical test. VAD samples exhibited an increase in alpha diversity compared to controls (p=0.072), indicating an increase in the diversity of bacterial communities found within VAD lung samples.. Beta diversity, or the measurement of diversity *between* samples,^64^ was performed on bacterial transcripts using the Bray-Curtis dissimilarity index (Figure 4B) and showed VAS and VAD cohorts clustered separately with a p-value of 0.01. These results provide evidence of a statistically significant shift in bacterial composition in VAD lungs at 8 weeks.

**Fig. 4.**
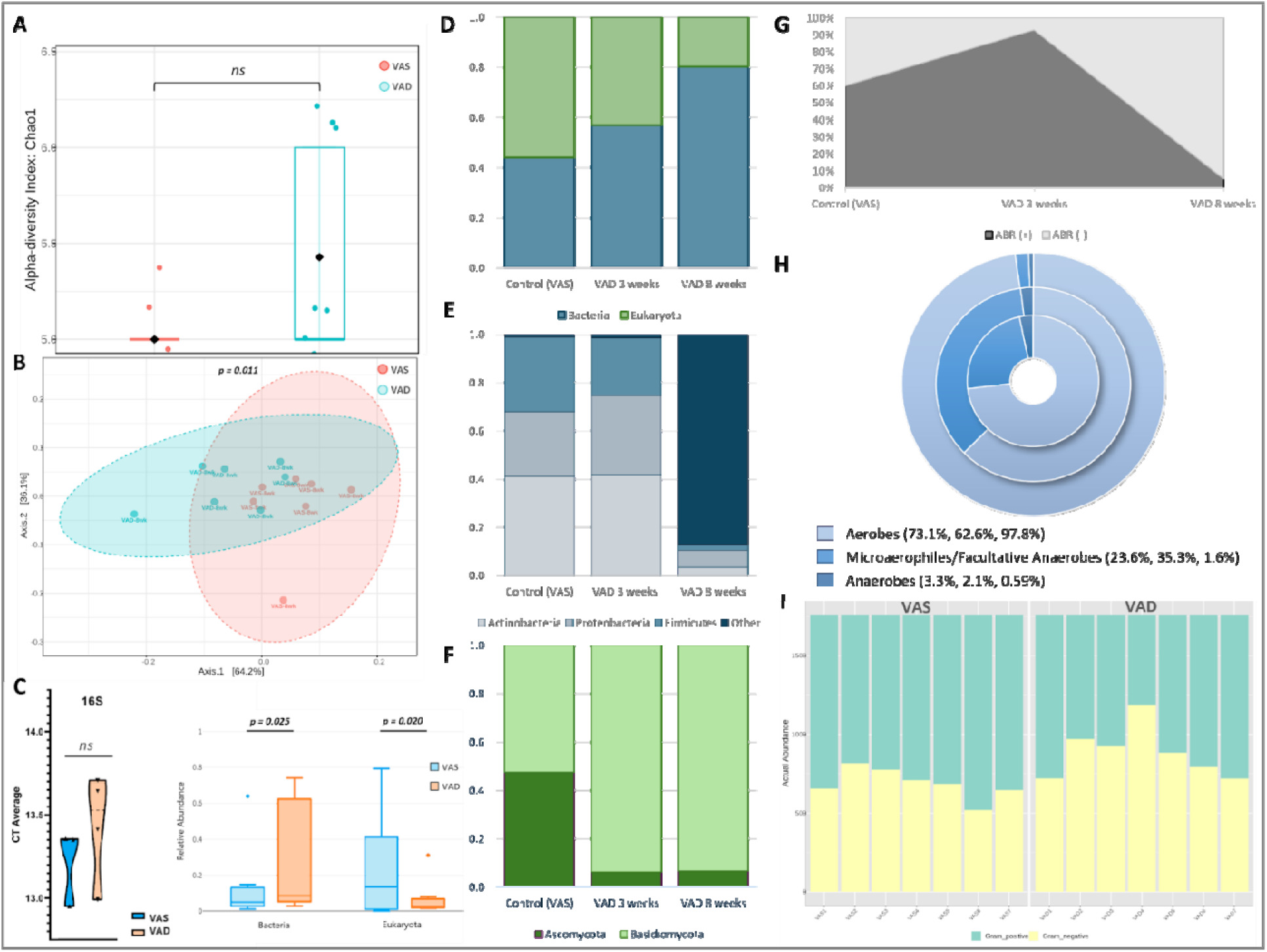
VAD alters microbial composition in the lower respiratory tract. (A) Alpha diversity barplot of bacterial transcripts at the Phylum level Item VAS and VAD lungs at 8 weeks using Mann-Whitney/Kruskal-Wallis statistical method, p=0.07 (n=14). (B) Beta diversity barplot of bacterial transcripts at 8 weeks using the 8ray-Curtis dissimilar^ index and PERMANOVA statistical method, p=0.01. (C) RT-PCR violin plot of 16S expression represented by CT Average (Y-axis, “СГ’ - cycle threshold) (left, n-S) and RNA-seq boxplots (right, n-14) of bacterial and fungal relative abundance In VAS and VAD lungs, p=0.02S and 0.020 respectively. (D-F) Relative abundance barplots of Bacteria and Fungi In VAS (n-18) and VAD lungs et 3 weeks (n-9) and S weeks (n-7). Y-axis represents the overall microbial abundance where 1.0 -100% оГ RNA transeripts. All bacteria with less than 1% abundance In VAS lungs were grouped Into ‘Other*, (ffl) Stacked proportion chart showing changes In relative abundance of antibiotic resistent (ABR+) and nan-ant biotic resistent (ABR-) bacteria In VAD lungs over time (n-32). (H) Slacked doughnut abundance plots of bacterial oxygen tolerance In VAS (Inner ring), VAD at week 3 (middle ring) and VAD at week 8 (outer ring) lungs, ft) Stacked barchart showing actual abundance of gram-positive end gram-negative bacterial transcripte at 8 weeks (n-14). Classifications of antibiotic resistance, oxygen tolerance, and gram status were manually annotated from BacDive.

To see if the observed shift translated to an increase in bacterial burden, we measured the 16S gene expression within VAS and VAD lungs using qPCR. Briefly, qPCR is a method of quantifying the number of sequence copies within a sample. By amplifying the bacterial 16S cDNA within each sample, we are able to quantify the total bacterial content and calculate the bacterial load within a group of samples.^65^ QPCR results from whole lung homogenates (n=8) did not show a significant change in bacterial burden (p = 0.42, Figure 4C, Table S1), but these findings do not fully explain the functional shift in the VAD lung microbiome over time.

While absolute abundance is often considered more biologically relevant, many microbiome researchers choose to calculate the relative abundance of microbes within a host tissue instead, as these values can be more functionally relevant to predict dysbiosis and disease progression over time; parametric models like DESeq2 uses negative binomial distributions applied to normalized data to minimize hidden confounding factors from sample fractions.^66^ By applying DESeq2 with FDR correction to our Log-transformed RNA read counts at 8 weeks (n=14), we confirmed both bacteria and fungi exhibited significant shifts in relative abundance (p = 0.025 and 0.020 respectively, Figure 4C). Specifically, relative bacterial burden did increase in VAD lungs over time (Figure 4D), although major shifts in bacterial composition were not evident until week 8 (Figure 4E) where pathobionts such as Planctomycetes were the most abundant.^67^ Meanwhile, the relative abundance of Ascomycota - a fungal phylum known to be beneficial for human health - plummeted as early as week 3 and maintained the same level of abundance at week 8 (Figure 4F).^68–70^

To determine the functional relevance of the compositional changes in VAD lungs over time, the bacterial metadatabase BacDive was used to annotate the RNA read counts of whole lung samples collected at 3 weeks and 8 weeks (n=32).^71^ VAS samples at 3 weeks and 8 weeks were combined to represent the control lung cohort (n=16), and any differences in the relative abundance of VAS microbes between these two time points at the Phylum level were insignificant (data not shown). Interestingly, the relative abundance of bacteria classified as antibiotic resistant are largely depleted at week 8 (Figure 4G). This change in gene expression could be due to the increased relative proportion of rare and low-abundance bacteria in VAD lungs that have yet to be classified as functionally relevant to human health. We also observed an overabundance of aerobic bacteria at the 8-week timepoint in the VAD lung (Figure 4H, outer ring). Since vitamin A is an antioxidant,^72^ these results suggest chronic VAD may lead to an increase in oxidative stress within the lung, which in turn could lead to the preferential selection of oxygen-tolerant microbes. We also observed an increase in the relative abundance of gram-negative bacteria compared to gram-positive bacteria in VAD lungs at 8 weeks, suggesting either a selective increase in gram-negative bacteria or transcriptional quiescence of gram-positive bacteria (Figure 4I).

Of note, our results are consistent with human studies where bacteria undergo a functional change towards colonization and oral biofilm formation; researchers investigating the development of dental plaque following a meal observed a reduction in the relative abundance of gram-positive bacteria in late biofilm samples, with different molecular pathways activated between early and late biofilm formation.^73^ Specifically, KEGG analysis of late biofilm transcripts were enriched for pathways involved in cellular transport, cell motility including pilus and flagella assembly, and DNA/mismatch repair, the same pathways enriched in our mouse dataset (Figures 3G-H, other data not shown). Therefore, our data suggests chronic VAD shifts the microbial composition of the LRT towards dysbiosis and early pathogenesis. This shift is evidenced by a reduction in the relative abundance of Ascomycota and gram-positive bacteria, as well as an increase in the relative abundance of oxygen-tolerant microbes more broadly, predisposing the VAD lung to infections and disease.

### VAD ENCOURAGES BACTERIAL COLONIZATION IN THE LOWER RESPIRATORY TRACT

To determine how VAD-induced lung dysbiosis influences the localization and colonization of bacteria within the LRT, we analyzed our 8-week RNA-seq samples using MicrobiomeAnalyst, a metacoder analysis package^76^ used to depict taxonomic differences between the VAS and VAD microbial communities.^77^ Our data shows that *Pseudomonadota* and *Deinococcota* are predominantly found in the VAD lung (blue), two Phyla known for their pathogenic and extremophilic potential (Figure 5A).^78,79^ In fact, *Deinococcota* is known for being able to synthesize its own carotenoids such as deinoxanthin, aiding its survival during hostile and nutrient-depleted circumstances.^79^ On the other hand, the commensal *Bacillota* Phylum is predominantly expressed in the VAS lung (red), suggesting that chronic VAD leads to a reduction in the relative abundance of commensal bacteria at the Phylum level.

**Fig. 5.**
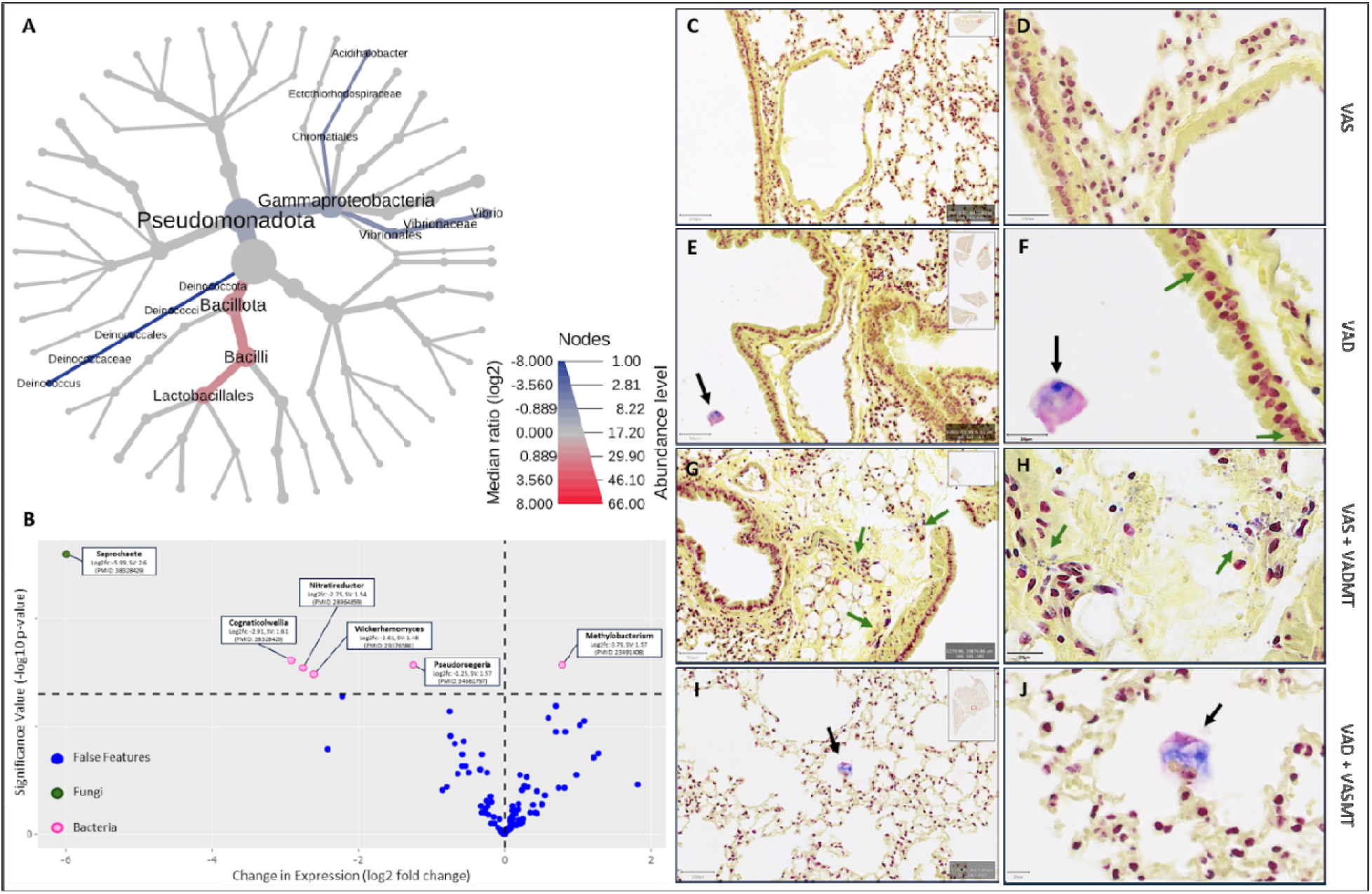
VAD encourages bacterial colonizatlon In Ille lower respiratory tract. (A) Heat tree or Bacterial OTU median ratios and relative abundance a: the genus level calculated using the Melacoder app in MicroolomeAnalyst. Blue = Predominantly In VAD (p < 0.05}. Red = Predominantly № VAS (p < O.C5). (B) Volcano pot or microbes differentially expressed In VAD lings at the species level using BatchQCversion 1.99.02 n Rstudlo version 4.3.1. (C.J) Bacterial gram stains of VAS (C,D) end VAD (E,F) mouse lungs at 8 weeks. (G.J) Gram stains of niratracheal microbial transplant recipients; VAS and *WD* lungs received VAD and VAS microbial transplants respectively for 3 weeks. Diet was maintained ’or’ 1 weeks overall Black arrows indicate 3-dimensional structures (oiofihls) containing 1;ram1losilive bacteria. Green arrows indicate mobile or bacteria that are still mobile or nonacherant along the airway.

To analyze microbiome data at the species level, we used the BatchQC^80^ Shiny App R package to correct our raw read counts for unidentified batch and covariate effects that could skew our results. Using the app, we first normalized the species-level log-transformed read counts using DESeq,^66^ followed by Limma^81^ for differential expression analysis (Figure 5B). Our volcano plot depicts the p-value (x-axis) and significance value (y-axis) for the compositional change of each species. Of the six microbes that met the significance threshold, Saprocaete from the Ascomycota Phylum was the only fungi significantly altered (green dot). We also observed an increase in Methylobacterium expression from the *Pseudamonadota* phylum, a species known for healthcare-associated infections.^82^

While changes in gene expression can indicate changes in microbial abundance, significant alterations in gene expression within a particular taxa can also indicate a functional shift from a *planktonic* or non-adherent state to a *sessile* or adherent state.^75^ To determine whether the VAD-induced dysbiosis translates to changes in lung colonization patterns, we analyzed bacterial Gram stains of the mouse airway after 8 weeks of dietary modulation (Figures 5C-F). We observed 3-dimensional biofilm structures containing gram-positive bacteria in VAD lungs (Figures 5E-F, black arrows) that were not observed in VAS lungs. The exception was a VAS sample with a biofilm attached to a vessel containing a granuloma (data not shown). We also observed a large cohort of planktonic gram-negative bacteria along the VAD airway epithelium (Figure 5E, green arrows), indicating that gram-positive and gram-negative bacteria may have distinct stress responses associated with host VA status in the lung.

To determine causality in relation to lung-microbe interactions, we wanted to investigate whether the microbiome can rescue or exacerbate the VAD and VAS lung phenotypes respectively. To that end, we extracted the lung microbiome from mice maintained on diets for 8 weeks and administered daily intratracheal (IT) injections of VAS and VAD microbes into VAD and VAS mouse lungs respectively for 3 weeks. It is important to note that while our extraction technique removed larger host and immune cells, signaling molecules and other immune cell milieu could not be removed from the samples using our extraction and IT protocol.^83^ Therefore, our daily microbial transplants contained the microbes and molecules found within the airway of the LRT. Interestingly, early signs of invasion and colonization by gram-positive bacteria were observed in VAS lungs exposed to VAD microbial transplants for 3 weeks (Figures 5G-H). VAD lungs receiving VAS microbial transplants also had evidence of biofilm formation (Figures 5I-J), although these structures could have persisted from the 8-week time point when the IT microbial transplants began.

These results confirm that host VAD status leads to bacterial invasion, colonization, and an increased risk of infection in the LRT of adult mouse lungs. While our transplant data provides evidence that VAD microbes exhibit post-biofilm invasion and colonization patterns in VAS lungs, more evidence is needed to determine whether a VAS microbial transplant is an effective prophylaxis towards slowing the progression of epithelial remodeling, dysbiosis, and bacterial colonization in VAD lungs.

## DISCUSSION

Microbial dysbiosis leads to increased inflammation, infections, and disease in the lung.^75^ For instance, significant alterations in the abundance of *Pseudamonadota* was observed in patients during COPD exacerbations.^74^ Our work provides evidence that dietary vitamin A is important for a healthy lung, while VAD alters the lung epithelium and the microenvironment to create a permissive niche for pathogenic outgrowth and biofilm formation.

Many papers have uncovered links between nutritional status of humans and the onset of lung cancer and disease, but there are currently no published studies indicating dietary VAD acts directly on the lung microbiome, nor is there evidence of host-microbe interactions in the lung that are mediated by host vitamin A status. This publication is the first to characterize the functional interactions between vitamin A metabolites, the lung, and the lung microbiome. Our lab has shown that VAD leads to epithelial remodeling and denudation in the airway, a loss of apical protein expression in the airway, focal hyperplasia, and a breakdown of mucociliary clearance, a critical mechanism responsible for lung homeostasis. These changes compromise the integrity of the epithelial barrier and create a permissive niche for pathogenic outgrowth and biofilm formation.

Metatranscriptomic analysis showed a reduction in fungal abundance and an increase in bacterial abundance in VAD lungs at week 8. Histological staining of bacteria confirmed these findings, while also providing evidence of a VA-mediated mechanism that triggers colonization in gram-positive but not gram-negative bacteria, suggesting that vitamin A may be involved in gram-specific quorum sensing mechanisms. Transcriptional data confirms changes in fungal composition precede measurable changes to the bacterial consortium, suggesting that VA-mediated quorum sensing mechanisms in the LRT are polymicrobial in nature.

A promising future direction of this work towards detection of respiratory infection is the observed fungal sensitivity to host vitamin A status. Ascomycota, the most abundant fungal Phylum found in humans worldwide, dramatically reduced transcriptional activity at our earliest time point in VAD lungs; this change in transcription preceded the hyperplasia and the bacterial biofilm formation. Therefore, future studies investigating the *mycobiome* and vitamin A may uncover useful and sensitive biomarkers for early detection of airway remodeling and infection in malnourished patients.

## CONCLUSION

Dietary vitamin A deficiency (VAD) causes epithelial remodeling in the lower respiratory tract characterized by epithelial denudation, focal hyperplasia, thickening of the smooth muscle, and a loss of ciliary function. Focal hyperplasia leads to an enrichment of secretory cells and a loss of apical secretory and ciliary protein expression in the airway, leading to a breakdown in MCC. These changes create a permissive niche for lung dysbiosis, where low-abundant and potentially pathogenic microbes are enriched and a functional shift from planktonic to sessile adhesion is observed in gram-positive bacteria, leading to biofilm formation.

## MATERIALS AND METHODS

### Housing and Dietary modulation

Wild-type C57BL/J6 female mice were purchased from Jackson Laboratory at 7-9 weeks of age (catalog # 000664). Female mice were used exclusively for the preliminary study to minimize in-fighting and potential injury during transit or during the required 1-week acclimation period following arrival to our animal facility, which would minimize the potential for a compromised immune system. Female mice also allowed us to co-house mice shipped in different cages, which would enable us to control for cage effects in our preliminary study.^84^ We also used female mice to remove sex effects from the preliminary results, which is another covariate that could have skewed our findings. Mice were purchased as littermates or siblings to minimize genetic drift as another covariate. Four mice were shipped in each cage; following the acclimation period, mice were separated evenly into each arm of the study and housed with a female from a different cage to ensure any variation in the microbiome due to cage effects was shared evenly between the two dietary treatments. Cage localization and variation in microbes on the cages themselves can drastically affect the outcome of lung microbiome research.^84^ Considering these findings, we changed the cage, food, water, and bedding as well as rotated the cages in the rack every 2-3 days so that each cage was never 1) next to the same cages or 2) in the same location on the rack more than once throughout the entire study.

We provided each cage with a custom research diet created by Envigo® that was either Vitamin A-sufficient (VAS) or VAD (Catalog # TD.91280 and TD.86143). The chow was sterilized, irradiated, and vacuum-sealed to prevent microbial contamination. The left lobe of each lung was paraffin embedded for histology and immunohistochemistry, while the right lobes were homogenized for mixed host-microbe bulk RNA-seq (Figure 1A).

### RNA Isolation and Extraction

The following protocol was designed by our lab to achieve mixed RNA samples with a higher microbiome RNA yield than previously published techniques. To do so, we tested many approaches and combined portions of three separate protocols for maximum results; a portion of our modified GentleMACS® technique created by our lab for FACS sorting and whole lung digest, a portion of the Qiashredder® tissue disruption and homogenization protocol to maximize mammalian RNA extraction, and a portion of the PureLink® RNA extraction and DNase Digest for maximum 16S and mixed microbiome sample RNA integrity. The following is our novel mixed host-microbe RNA extraction protocol:

#### Preparation

Mice were euthanized using isoflurane with cervical dislocation as a secondary method. To ensure minimal cross-contamination, dissection tools were sterilized in 100% ethanol for one hour before placing in a surgical tray. Dissection supplies and pre-aliquoted sterile solutions were placed in a metal surgical tray, the tray sealed in a transparent bag, and processed through a ThermoFisher CL-1000 UV crosslinker once (catalog # 95-0174-01, serial # 061608-001). Without opening the bag, items were turned over and processed through the UV crosslinker again. The supplies remained in the sealed transparent bag until it was time for lung extraction. The bag was sterilized with 70% ethanol before placing in a fume hood to open for use.

#### Extraction

Lungs were extracted using sterile forceps and dissecting scissors. The trachea and as much of the connective tissue as possible were removed from the lower respiratory tract (LRT) while the sample was still within the thoracic cavity to minimize contamination. The left lobe was placed in a canonical tube containing 4% paraformaldehyde and placed on ice. The right lobe was placed in a 1.5mL Eppendorf tube, previously sterilized in the surgical tray, containing 1mL of RNAlater and placed on ice. The left lobes were stored in a 4°C refrigerator until the next day.

#### Whole Lung Tissue Disruption and Homogenization

Each lobe that was previously placed in RNAlater was transferred to a gentleMACS® M-tube with 1.2mL of RLT buffer from the Qiagen® RNeasy mini kit. 24_µ_L 2M DTT was added to the tube, and then homogenized using the gentleMACS^TM^ Octo-dissociator). Two programs were used that are pre-programmed and standard for this machine; the “m_lung_01_01” setting was run once, followed by two rounds of “RNA_01_01”. The M-tubes were then centrifuged on 15,000 rpm for 2 minutes. Up to 650 _µ_L of the supernatant was transferred into a QiaShredder spin column by Qiagen® (catalog # 79656), keeping the M tubes on ice. Qiashredder columns were centrifuged at 12,000 xg for 2 minutes before adding 50 _µ_L of RNA-free water directly onto the spin column and centrifuging for another 2 minutes at 12,000 x g. The flow-through tubes were capped and placed on ice between each transfer of the spin column.

#### Pellet Disruption and Homogenization

100 _µ_L of lysozyme solution was added to the M tubes (see STARS methods section) and vortexed to resuspend the pellet. 8 _µ_L of 1% SDS solution was added before vortexing and transferring the samples to Eppendorf tubes. Using a 21-gauge needle, we rapidly filled and expelled a 1 mL syringe 15-20 times to ensure the bacterial cell walls had been denatured. The tubes were then centrifuged at 12,000 x g for 2 minutes at room temperature before transferring the supernatant to a clean RNA-free microcentrifuge tube.

#### RNA Extraction

250 _µ_L of 100% ethanol was added to each tube and vortexed briefly. Each sample was transferred into a spin cartridge with a collection tube from the Purelink® RNA extraction kit and centrifuged at 12,000 x g for 15 seconds at room temperature. The flow-through was discarded but not the collection tube. 350 _µ_L of Wash Buffer I was added to the column to repeat the previous step once before discarding and replacing the collection tube. We were sure to use only one spin cartridge per sample tube collected from the homogenization steps. Even though fewer spin cartridges could reduce contamination and the potential for sampling bias, performing too many centrifuges with the same spin cartridge will clog the membrane and dramatically reduce the RNA yield.

#### DNA Digest

80 _µ_L of DNase Digest solution from the Purelink® DNase digest protocol was added to each spin cartridge and incubated for 15 minutes at room temperature. 500 _µ_L of Wash buffer II was added to each spin column and centrifuged for 15 seconds at 12,000 x g. Flow-through was discarded from the collection tube before repeating the previous step. The spin cartridge was placed into a clean 1.5mL Eppendorf tube for its final elution and collection phase. 40 _µ_L of RNA-free water was added to the <u>center</u> of the spin column before incubating for 1 minute followed by centrifugation at 12,000 x g for 1 minute. Another 40 _µ_L of RNA-free water was added around the <u>edges</u> of the spin column and the previous incubation and spin steps were repeated once per spin cartridge. Each spin column collected from the same sample was placed into the same Eppendorf collection tube for the elution stage. Each collection tube was vortexed to combine elutions well before verifying the RNA yield and quality using a Nanodrop^TM^ spectrophotometer.

Using the protocol outlined above, the RNA we collect averages a perfect nucleic acid 260/280 ratio of 2.0-2.1, and an excellent RNA Integrity Number (RIN) of 9.1-9.9. Our designed protocol is exhaustive, requiring an entire day for extraction and three separate RNA processing kits plus additional reagents. However, our downstream analysis results indicate these extra steps lead to higher yields and RINs than other protocols we’ve tried.

### Host-Microbe Total RNA Sequencing

#### Sample selection and sequencing depth

To ensure the best quality data, we extracted RNA from 18 samples after 3 weeks (9 VAS and 9 VAD) and 23 samples after 8 weeks of dietary modulation (11 VAS and 12 VAD) before submitting them to the Boston University Microarray and Sequencing Resource (BUMSR) Core Facility for QC analysis. The 10 samples at 3 weeks and 14 samples at 8 weeks with the best quality and yields were approved for further processing and library preparation, with 260/280 scores ranging from 2.0 to 2.1 and yields from 40.4 ng/uL to 284.6 ng/uL. To increase the sequencing depth of the microbiome in our mixed samples, BUMSR performed mouse rRNA depletion prior to sequencing. Next Generation total RNA-sequencing was performed using the Illumina NextSeq2000. Samples underwent two sequencing runs using a P2 flow cell for the first run and a P3 flow cell for the second run to increase sequencing depth. Each run repeated for 200 cycles with 100×100 base pair paired-end reads. The final sequencing depth was approximately 1.77 billion read pairs, ranging from 60.7-86.9 million reads for each of the 24 samples collected (**Table 3**).

**Table 1.**
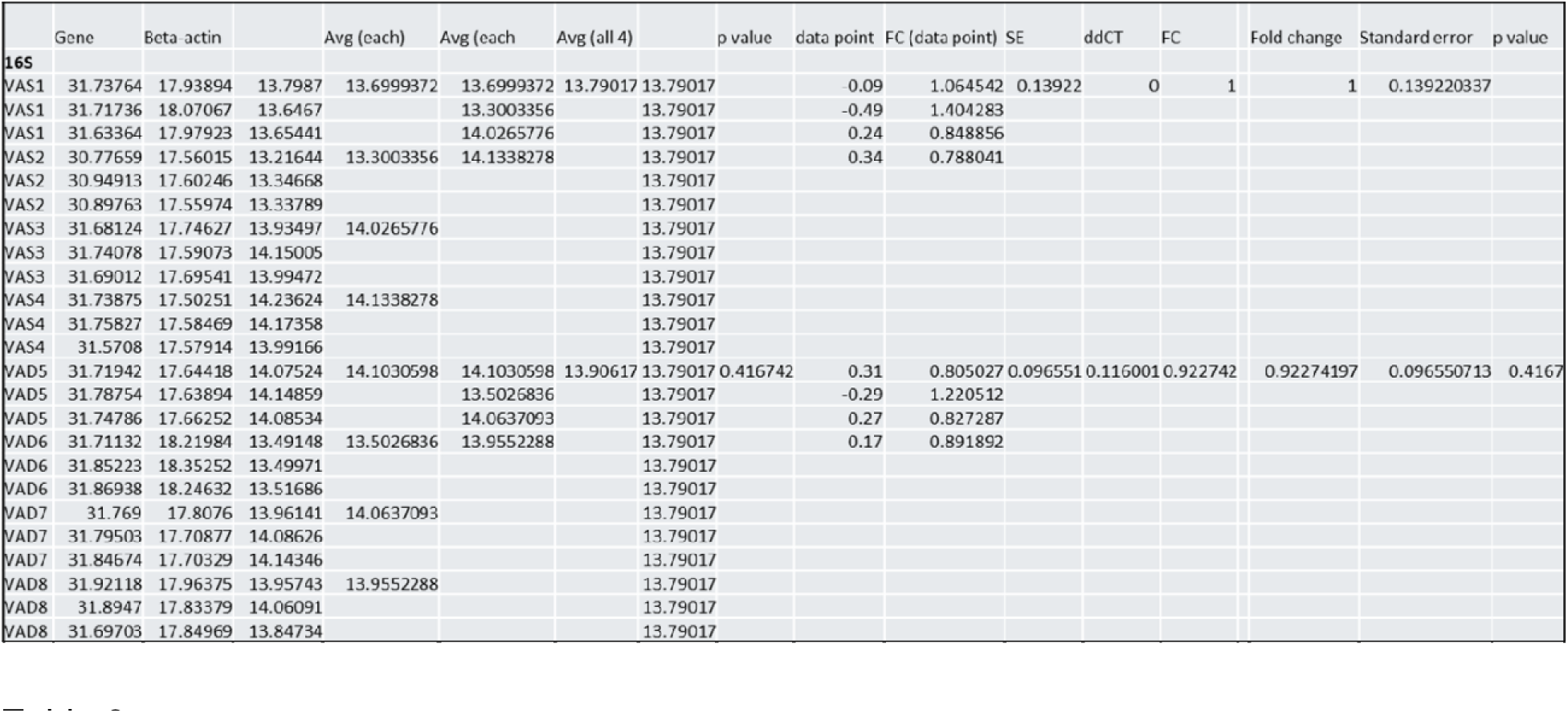

**Table 2.**
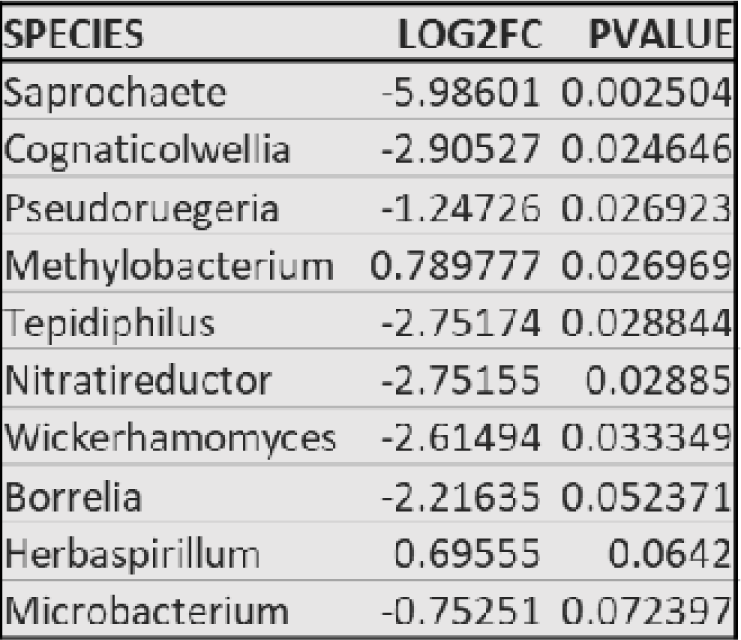

**Table 3.**

| Sequencing Summary |  |  | All Reads in Millions |  |  |
| --- | --- | --- | --- | --- | --- |
| Sample Name | Reads Obtained P2 Run (M) | Reads Obtained P3 Run (M) | Total Reads Obtained (M) | Metric | Reads (M) |
| VAS1_3wk | 17.76 | 49.86 | 67.62 | Minimum | 60.70 |
| VAS2_3wk | 17.71 | 49.56 | 67.27 | Maximum | 86.86 |
| VAS3_3wk | 15.67 | 45.03 | 60.70 | Mean | 73.97 |
| VAS4_3wk | 21.48 | 57.79 | 79.28 | Total | 1775.21 |
| VAS5_3wk | 20.47 | 54.24 | 74.71 |  |  |
| VAD1_3wk | 18.80 | 49.41 | 68.21 |  |  |
| VAD2_3wk | 16.18 | 46.50 | 62.67 |  |  |
| VAD3_3wk | 21.88 | 55.47 | 77.35 |  |  |
| VAD4_3wk | 16.47 | 45.36 | 61.83 |  |  |
| VAD5_3wk | 17.55 | 49.17 | 66.72 |  |  |
| VAS2_8wk | 18.44 | 49.34 | 67.78 |  |  |
| VAS3_8wk | 21.80 | 55.78 | 77.58 |  |  |
| VAS4-8WK | 23.20 | 57.56 | 80.76 |  |  |
| VAS5-8WK | 25.37 | 60.77 | 86.14 |  |  |
| VAS7-8WK | 20.04 | 55.84 | 75.87 |  |  |
| VAS9-8WK | 25.98 | 60.88 | 86.86 |  |  |
| VAS10-8WK | 19.34 | 50.81 | 70.15 |  |  |
| VAD2-8WK | 22.35 | 55.97 | 78.33 |  |  |
| VAD3-8WK | 23.03 | 57.61 | 80.64 |  |  |
| VAD4-8WK | 21.07 | 53.40 | 74.47 |  |  |
| VAD5-8WK | 19.67 | 54.70 | 74.37 |  |  |
| VAD7-8WK | 23.42 | 57.08 | 80.49 |  |  |
| VAD8_8wk | 21.94 | 55.37 | 77.31 |  |  |
| VAD10_8wk | 23.68 | 54.41 | 78.09 |  |  |

#### Sequence Pre-processing and Alignment

Raw FastQ files of single paired-end reads were trimmed based on minimum Illumina phred scores of 33 using the Trimmomatic package version 0.36.^85^ The trimmed sequences were aligned to bacterial, fungal, viral, and miscellaneous microbial gene indices downloaded from Refseq^86^ while filtering out samples that also aligned to human and mouse gene indices from Refseq. Rsubread was used to align the same sequences to the mouse gene index mm10.^87^ Raw RNA sequences feature counts and the pre-processing and sequence alignment code written by our lab is available on our Github repository.^88^.

### RNA-Seq Data Analysis

Raw read counts were loaded into two R-based Shiny Apps for preliminary data analysis: for the microbiome, Animalcules^89^ version 1.4.0 in RStudio version 4.0.2, and for the host, Single Cell Toolkit^90^ version 1.9.1 in RStudio version 4.1.1. Within the Single Cell Toolkit platform, the host RNA-seq read counts were normalized using the Limma R/Bioconductor software package,^91^ version 3.44.3 in Animalcules and version 3.48.3 in Single Cell Toolkit. Alpha and beta diversity plots were generated in MicrobiomeAnalyst.^76^ P-values for diversity plots were calculated in the app using the Wilcoxon Rank Sum Test with continuity correction model. BatchQC was used to normalize feature counts using Combat and DESeq.

### Paraffin-embedding and sectioning lung tissue

Left lobes were fixed in a 4% paraformaldehyde solution overnight at 4°C. We then paraffin-embedded the lungs by clearing the tissues in serial dilutions of ethanol and xylene and processing the tissues in a 60°C vacuum oven with paraffin wax for 3 hours. The tissues were left to harden at room temperature before storing in the 4°C fridge until sectioning. Paraffin-embedded tissues were sectioned into 5_µ_m tissue slices using a microtome. The tissues were left in a 37°C oven overnight to cure before storing in a 4°C fridge until staining.

### Immunohistochemical staining

Protocols for each stain were followed using the guidelines specified by their respective manufacturers; Scgb1a1/CC10 from Santa Cruz Biotech (catalog # sc-365992), Acetylated tubulin from Sigma-Aldrich catalog # T7451, and Foxj1 from Thermo Fisher Scientific catalog #14-9965-82. Our lab added 0.2% Tween at the blocking stage to permeate cells for sufficient surfactant protein C staining but all other staining protocols did not require this step. To assess changes in epithelial protein expression, we used immunohistochemical stains for CC10 (secretory), Tubb4a and acetylated tubulin (ciliary), and surfactant protein C (type I pneumocytes), although the majority of structural and functional changes explores focused on the ciliated epithelial cells.

### Ciliary Function Assay

Tissue collection, processing, and imaging for the ex-vivo tracheal analysis of ciliary motility is published and publicly available by Jove.^57^ We used a Nikon wide-field deconvolution epifluorescent microscope at the Boston University Medical Campus Cellular Imaging Core Facility. We used 0.18 m Fluoresbrite® carboxylate microspheres from Polysciences, catalog number 09834-10. Video footage of microsphere motility was recorded at 250 fps in accordance with the above mentioned Jove protocol. The videos of equal time length were then processed in the Nikon NIS Viewer package and composite images of each video file were generated to measure distance traveled by each bead using Fiji/ImageJ software.^92^

## ACKNOWLEDGEMENTS

We would like to acknowledge all the helpful feedback and support by the dissertation advisory committee: Bob Varelas PhD (chair), Melisa Osborne PhD, and Peggy Lai MD. Video microscopy was performed at the BU Imaging Core Facility. RNA sequencing was performed by the BU Microarray and Sequencing Resource Core Facility. Special thanks to Yuriy O.Alekseyev, Ashley Leclerc, Nickolas Gorham, and Christopher Williams for their help with optimizing our samples for bulk sequencing.

## FUNDING

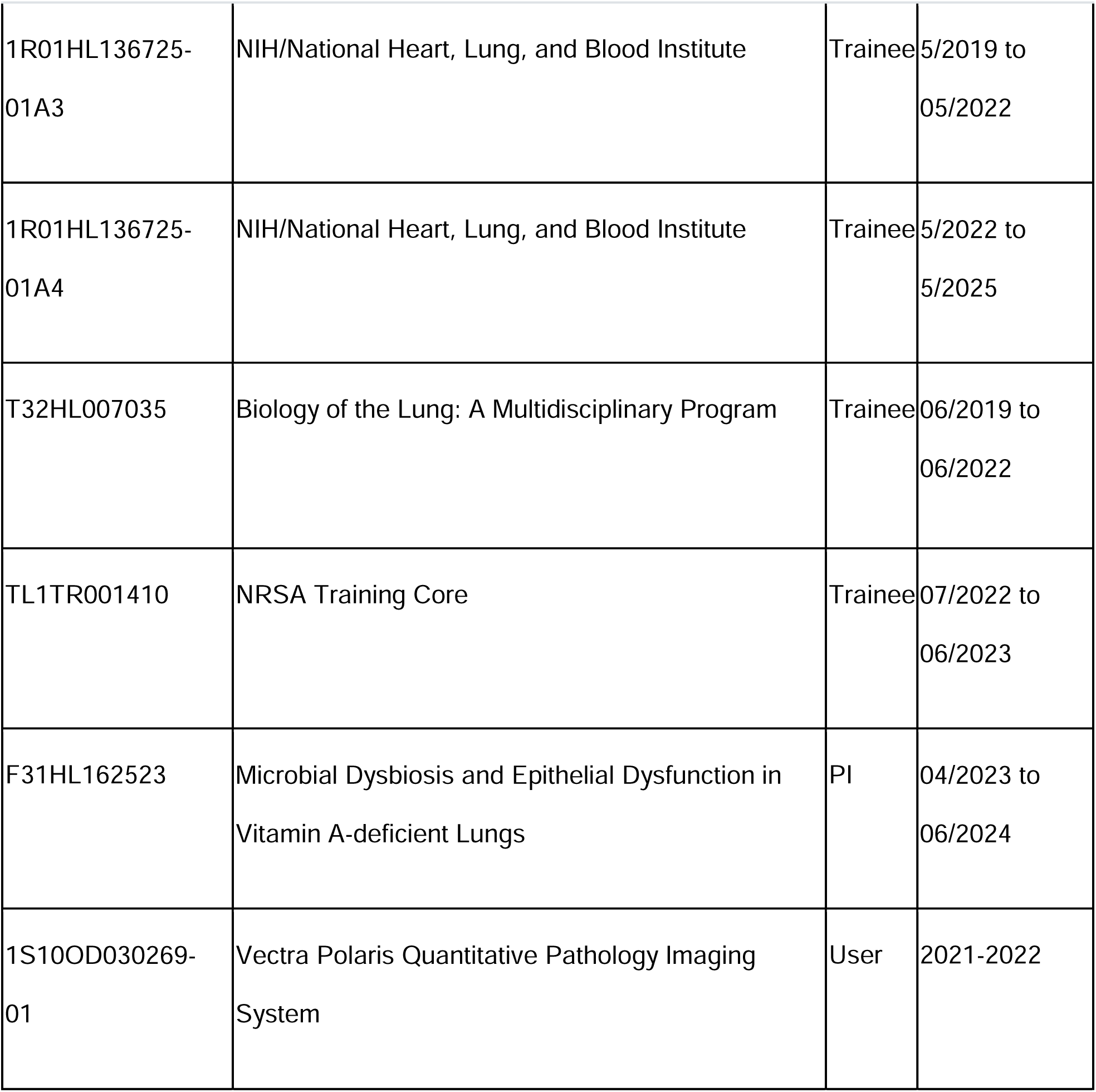

## AUTHOR CONTRIBUTIONS

Conceptualization: KQ, FC

Methodology: KQ, FC, WEJ

Investigation: KQ, FC, WEJ

Visualization: KQ, FC

Funding acquisition: KQ, FC

Project administration: KQ

Supervision: FC, WEJ

Writing – original draft: KQ

Writing – review & editing: FC, WEJ

## DATA AND MATERIALS AVAILABLE

https://github.com/Kiloni/VADandTheLungMicrobiome

